# Healthy Aging as Information Divergence in the Multiplex Brain

**DOI:** 10.64898/2026.05.29.728670

**Authors:** Debika Ghosh, Lucina Q. Uddin, Dipanjan Ray, Moumita Das

## Abstract

Understanding the co-evolution of the human brain’s structural scaffold and functional traffic across the adult lifespan remains a fundamental challenge in neuroscience. While age-related degradation in grey matter and functional activation is well-documented, the joint trajectory of the structural (SC) and functional (FC) connectomes is often overlooked due to the lack of an integrative framework. Here, we model the brain as a multiplex network to quantify the information-theoretic interdependencies between these two layers in a cross-sectional cohort of 589 healthy individuals (ages 18–88) from the Cambridge Centre for Ageing and Neuroscience (Cam-CAN) dataset. Using Jensen-Shannon Divergence and relative entropy metrics, we identify a fundamental organizing principle of healthy aging characterized by: a progressive information divergence where functional dynamics increasingly “untether” from their underlying structural constraints.

Our results reveal that this decoupling follows a robust linear trajectory, yet is highly spatially heterogeneous. Meso-scale community analysis using the Multiplex Map Equation identifies subcortical hubs—specifically the putamen, pallidum, caudate, and thalamus—as the primary epicenters of age-related SC-FC divergence. This topological shift toward functional independence in subcortical “switchboards” provides a mechanistic connectomic signature for the well-documented decline in fluid intelligence and motor adaptation that accompanies aging. In striking contrast, the limbic core (hippocampus and entorhinal cortex) exhibits remarkable stability across the lifespan, suggesting a biological imperative to preserve high-fidelity memory circuits amidst global communicative rewiring. By framing healthy aging as a systematic subcortical untethering alongside limbic resilience, our work provides a powerful new multiplex baseline to distinguish normative cognitive decline from the early topological signals of neurodegenerative disease.

**Significance Statement:** Healthy aging involves a fundamental reorganization of the brain’s information architecture. By modeling the brain as a multiplex network and leveraging information-theoretic measures, we demonstrate that adult lifespan development is characterized by a progressive “untethering” of functional dynamics from their underlying structural constraints. This decoupling is driven by a striking geometric paradox within subcortical hubs: while physical structural scaffolds undergo severe network constriction and isolation, their functional boundaries expand and blur in a state of neural de-differentiation. This architectural mismatch is mediated by a synchronized prefrontal-to-limbic pathway shift that operates collinearly with chronological age and mirrors the selective decline of fluid intelligence and motor adaptation that accompanies aging. Our work establishes a rigorous multiplex framework that moves beyond descriptive accounts of neural tissue loss to provide a normative baseline for distinguishing healthy brain aging from neurodegeneration.

## 1 Introduction

The quest to map the neural landscape of healthy aging is one of the most significant open questions in contemporary neuroscience. As the global population ages He et al. [2016], identifying the organizing principles that govern neurobiological transitions is essential for distinguishing normative maturation from the early stages of cognitive decline.

Historically, our understanding of the aging connectome has relied on a fragmented approach, examining structural Lo et al. [2011], Liu et al. [2017], Neudorf et al. [2024] and functional changes Edde et al. [2021], Doval et al. [2024] as distinct, isolated phenomena. While we have gained vital insights into white-matter degradation Mousley et al. [2025] and functional network reorganization Chan et al. [2014], Wang et al. [2012], this isolationist paradigm overlooks the intrinsically coupled nature of brain architecture. Studying these modalities in isolation fails to explain how functional neural traffic navigates a changing physical scaffold. To capture the true complexity of the aging brain, we must investigate the joint trajectory of the structural and functional connectome as a unified, interdependent system.

Characterizing these adult lifespan trajectories is essential to resolving a profound biological paradox at the heart of cognitive neuroscience: how does the functional brain maintain, and often reorganize, its network integration while its underlying physical white-matter infrastructure is progressively eroding? To reconcile this tension, a framework capable of simultaneously tracking anatomical constraints and functional traffic across time is strictly required. By employing a multiplex network framework to represent the structural and functional layers as a cohesive biological architecture, we move beyond simple single-layer correlations to uncover how the dependencies between modalities evolve with age. Unlike edge-wise correlational coupling metrics — which assume a linear correspondence between modalities, yield a single localized estimate, and are sensitive to parcellation and thresholding choices — information-theoretic measures quantify the full statistical distance between the entropy spectra of the structural and functional graph Laplacians, making them distribution-free, robust to differences in the sign and scale of structural versus functional edge weights, and sensitive to higher-order topological reorganization that correlational approaches cannot detect. Leveraging an information-theoretic approach, we utilize metrics such as Jensen-Shannon Divergence and Relative Information Loss to rigorously quantify the topological “distance” between the brain’s structural scaffold and its functional dynamics, providing a mathematical lens to evaluate exactly how strictly functional communication is governed by its anatomical wiring across the adult lifespan.

Studies of typical development in children and adolescents have demonstrated that structure-function coupling increases with age, particularly in brain regions supporting cognitive maturation, and that these developmental changes in SC-FC relationships are linked with individual differences in intelligence and executive function (Soman et al. [2023], Baum et al. [2020], Feng et al. [2024]). This evidence establishes SC-FC coupling as a meaningful marker of neurocognitive maturation earlier in the lifespan, but whether this coupling continues to strengthen, plateaus, or instead reverses across adulthood and older age remains unresolved. Notably, this developmental literature has largely characterized SC-FC coupling using regional or edge-wise correlational approaches, in which the structural and functional connectomes are computed separately and related post hoc through a single correlation coefficient per region or edge. While informative, such approaches cannot capture higher-order interdependencies between the two modalities as a unified network system, and provide only a localized rather than system-wide description of structure-function correspondence. Characterizing the normative trajectory of SC-FC relationships in healthy aging using a framework that instead treats structure and function as a single interdependent system is therefore essential — not only to determine whether the coupling gains established in youth persist into later life, but to identify which circuits maintain versus lose this alignment, and how this distinction relates to resilience or vulnerability in cognitive function among older individuals.

Drawing on a cross-sectional cohort of 589 healthy adults from the Cam-CAN dataset (Shafto et al. [2014], Taylor et al. [2017]), we propose a fundamental organizing principle of healthy aging: a progressive *information divergence* between structural and functional networks. Our analysis reveals a continuous, linear “untethering” across the adult lifespan, where functional dynamics increasingly drift apart from their deterministic physical wiring. We conceptualize this divergence as a bio-evolutionary trade-off, where the brain transitions from a state of youthful structural efficiency to an older state of functional degeneracy.

Crucially, we show that this decoupling is not spatially uniform; rather, it is uniquely localized to subcortical hubs—the brain’s central “switchboards,” including the putamen, pallidum, caudate, and thalamus. By resolving this node-level untethering at both the architectural and pathway scales, we reveal that this divergence is driven by a fundamental structural-functional geometric mismatch. While the physical structural infrastructure undergoes severe, synchronized network constriction that isolates these subcortical epicenters, their corresponding functional communities completely resist this collapse, instead expanding and smearing their boundaries into unconstrained cross-talk. We demonstrate that this geometric crossover is mediated by a highly coordinated, edge-wise pathway migration across the lifespan, where both layers progressively sever their ties with the executive prefrontal cortex to hyper-consolidate within the primitive limbic core, albeit at radically mismatched layer velocities. While this functional expansion and pathway rerouting may represent a dynamic compensatory attempt to preserve network traffic over an eroding physical skeleton, our behavioral models demonstrate that it operates collinearly with chronological aging, acting as a direct proxy for the degradation of fluid intelligence and motor adaptation. In striking contrast, the primary limbic core acts as a resilient anchor, maintaining rigid structural-functional alignment even as global communication rewires, potentially to protect core memory networks from catastrophic representational corruption.

By quantifying this structural-functional drift, this work moves beyond descriptive accounts of neural decline, providing a new mechanistic framework for understanding the physical limits of cognitive resilience, reorganization, and fluid intelligence across the human lifespan.

A schematic illustration of our analytical pipeline is provided in Figure 1.

**Figure 1:**
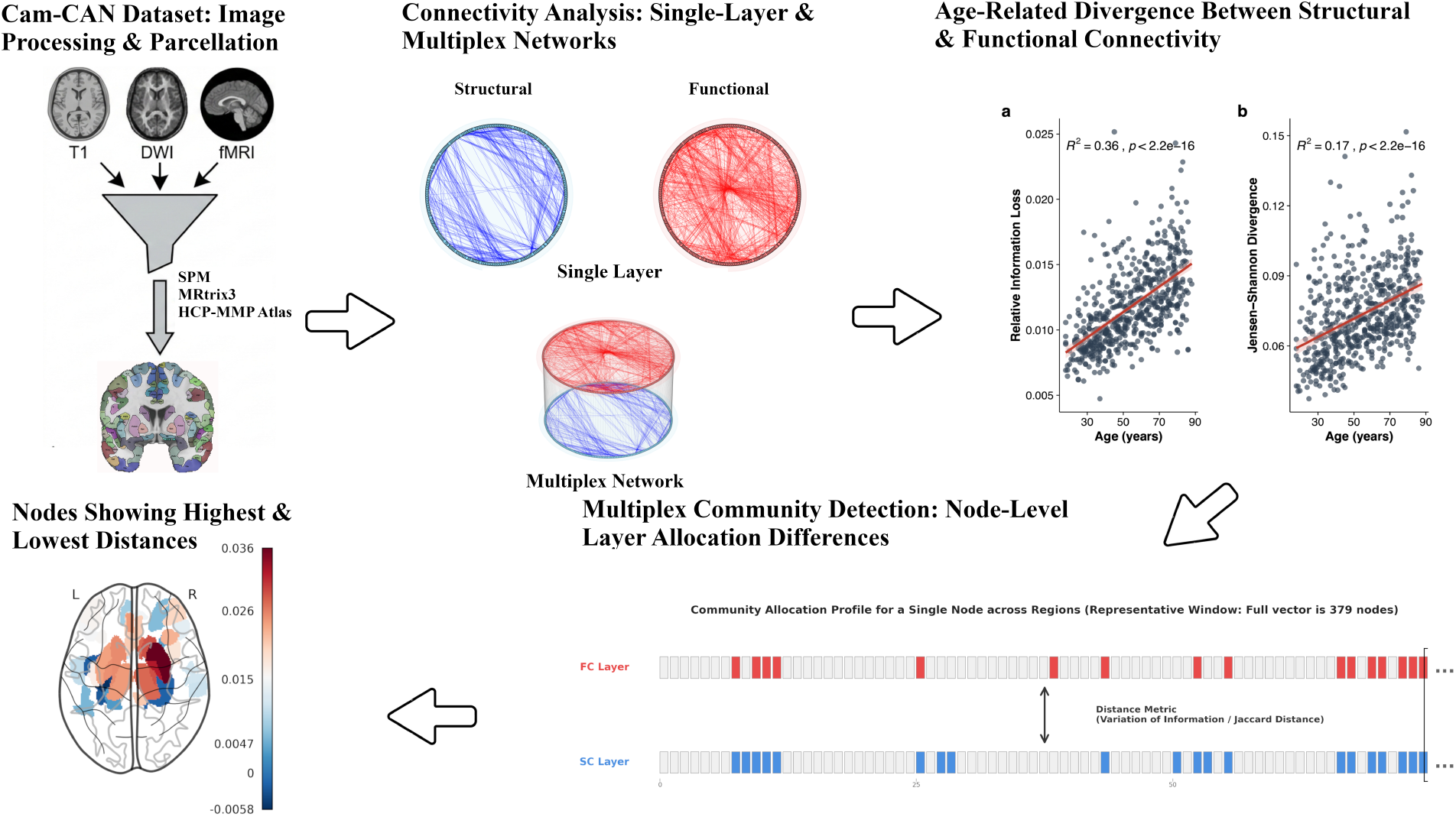
Analysis Pipeline. The schematic illustrates the extraction of structural and functional connectomes, the construction of the multiplex network, and the subsequent information-theoretic evaluation of structural-functional divergence across the lifespan.

## 2 Results

To characterize the trajectory of structural-functional (SC-FC) coupling across the adult lifespan, we constructed and analyzed multiplex connectomes for a cross-sectional cohort of 589 healthy adults (aged 18–88). Subjects were stratified into 12 age groups to assess lifespan trajectories. The multiplex analyses were performed using the muxViz framework De Domenico et al. [2015a], De Domenico [2022].

### Global Structural-Functional Information Divergence

We first investigated whether the functional dynamics of the aging brain become increasingly “untethered” from their underlying structural constraints. To test this, we evaluated the relative information loss (reducibility, *q*(*C*)) and the Jensen-Shannon Divergence (*d*_JS_) between the SC and FC layers across the lifespan.

Our analysis reveals a progressive, global information divergence with age (Fig. 2). The relative information lost when collapsing the multiplex layers into a single aggregated network (*q*(*C*)) systematically increased across the lifespan, rising from approximately 1% at age 18 to nearly 2% by age 89 based on model predictions. A linear regression confirmed a significant positive association between age and multiplex reducibility (*β* = 9.7 *×* 10^−5^, *p <* 0.05), with age explaining a substantial portion of the variance (*R*^2^ = 0.358).

**Figure 2:**
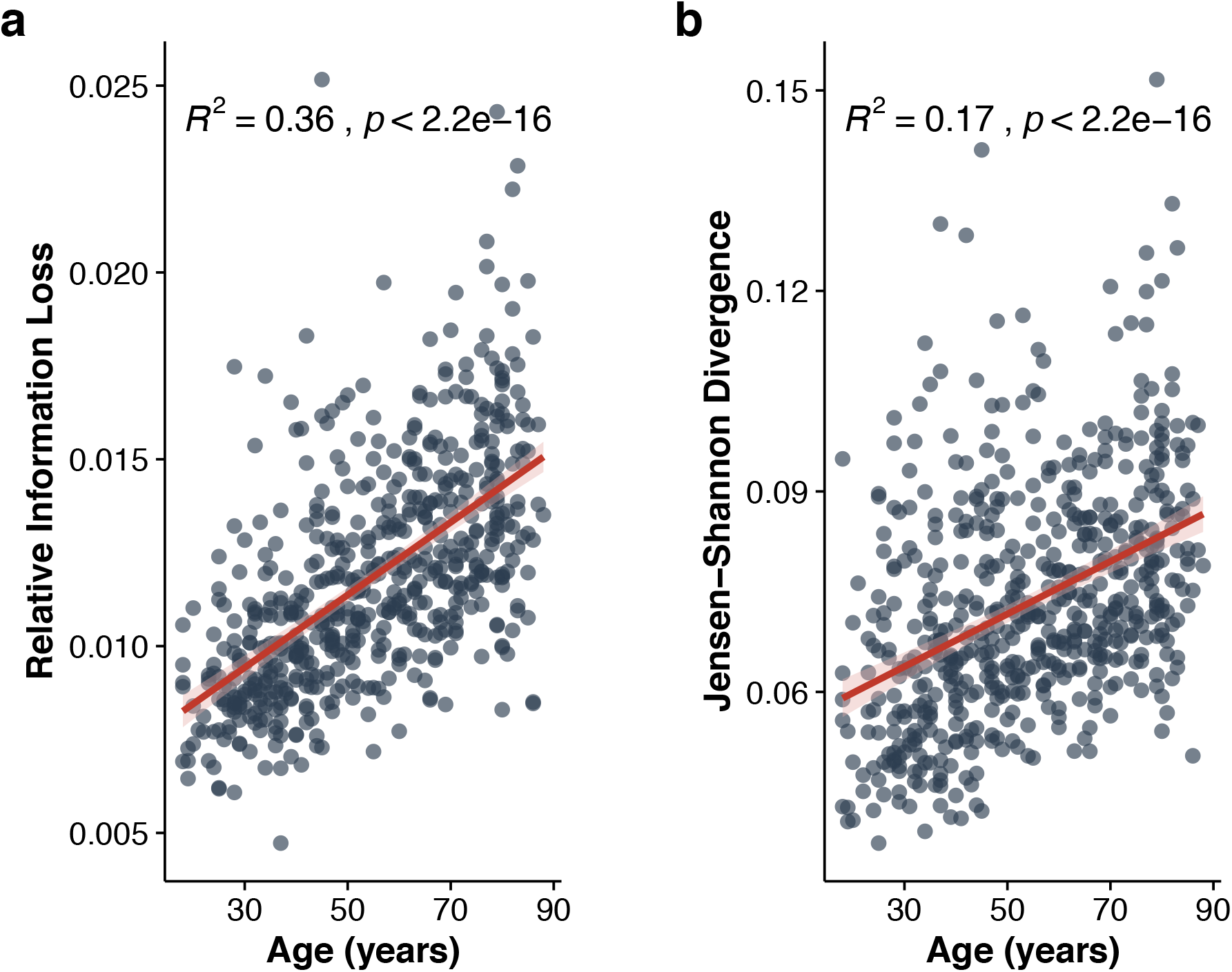
Global Multiplex Information Divergence Across the Adult Lifespan. **(a)** Relative Information Loss (*q*(*C*)) derived via Von Neumann entropy increases progressively across age groups (*R*^2^ = 0.36, *p <* 2.2 *×* 10^−16^). **(b)** Jensen-Shannon Divergence (*d*_JS_) increases linearly with chronological age (*R*^2^ = 0.17, *p <* 2.2 *×* 10^−16^).

This growing necessity of the multiplex representation is driven by an increasing dissimilarity between the physical and functional topologies. The Jensen-Shannon divergence (*d*_JS_) exhibited a parallel age-related increase, shifting continuously from ~ 5% in early adulthood to ~ 8% in later life. This linear increase was highly significant (*β* = 3.9 *×* 10^−4^, *p <* 0.05, *R*^2^ = 0.172), quantitatively demonstrating that as the brain ages, its functional traffic operates with increasing independence from its deterministic structural scaffolding.

### Subcortical Hubs are the Epicenters of Age-Related Decoupling

To pinpoint the anatomical drivers of this divergence, we quantified the node-level mismatch between structural and functional community assignments. We fitted a linear mixed-effects model using the normalized Variation of Information (VI) as the outcome metric, incorporating subject-specific and node-specific random effects to account for baseline individual and regional variability.

At the mean age of the cohort, the baseline normalized VI between SC and FC community profiles was substantial (mean = 0.71, SE = 0.009). Crucially, age was a highly significant positive predictor of further community mismatch (*β* = 0.063, SE = 0.0082, *p <* 0.05), confirming that the spatial allegiance of brain regions across structural and functional layers drifts apart over the lifespan. Node-specific random slopes indicated considerable anatomical heterogeneity in this aging effect (SD = 0.013).

To identify which specific regions drive this untethering, we evaluated the age-related trajectory of each node, applying a False Discovery Rate (FDR) correction (*q <* 0.05). The most profound structural-functional decoupling was localized to subcortical hubs (Fig. 3). Specifically, the bilateral **putamen, pallidum, caudate, and thalamus**—alongside a localized subset of cortical nodes—exhibited the steepest age-related increases in both Variation of Information and Jaccard distance. These findings indicate that the brain’s subcortical “switchboards” are the primary epicenters of the aging process, transitioning toward a state of heightened functional independence from their physical white-matter wiring.

**Figure 3:**
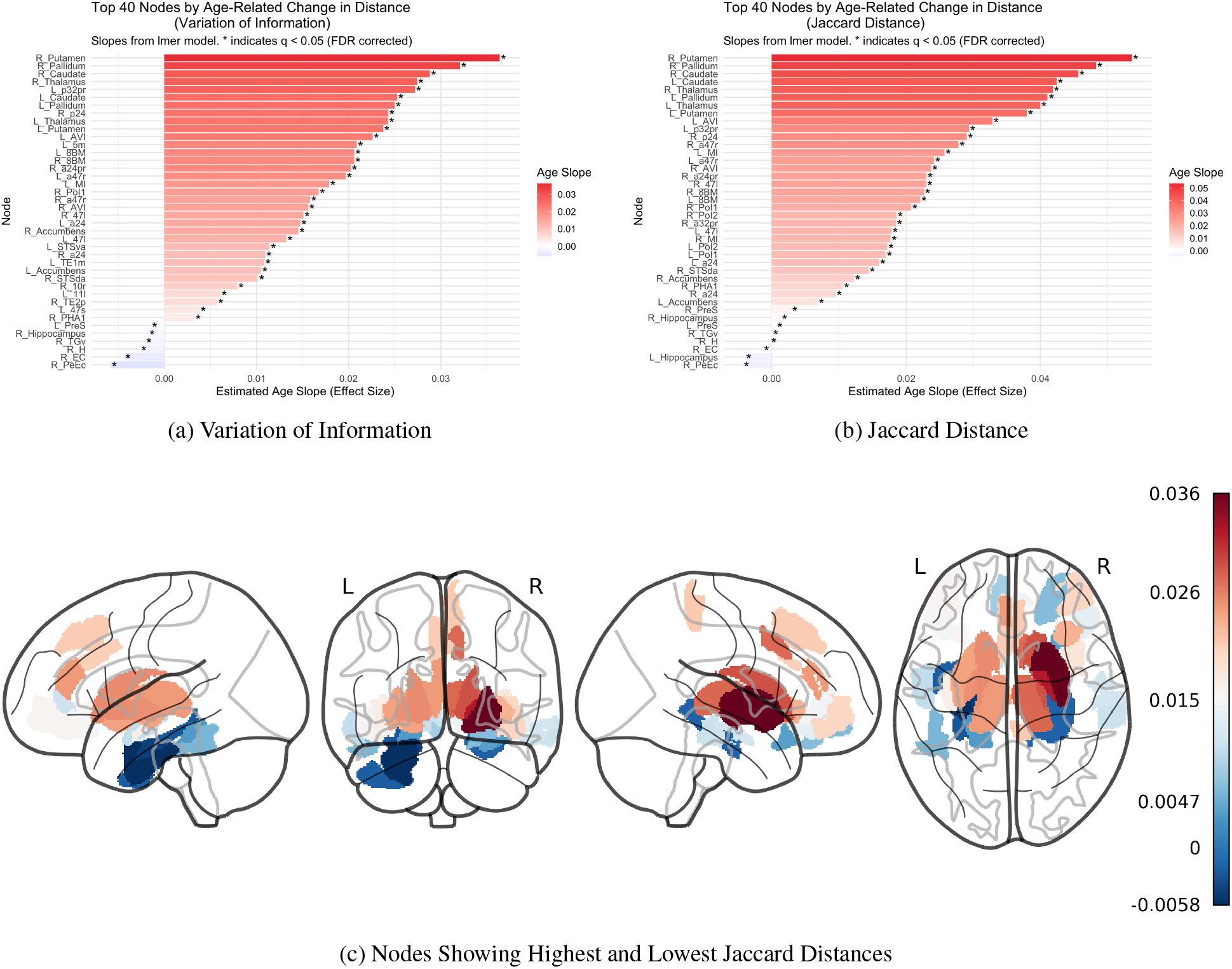
Regional Heterogeneity of Age-Related Structure-Function Reorganization. Node-level mapping of topological changes derived from the Multiplex Map Equation (*r* = 0.15) across the adult lifespan (*N* = 589). **(a–b)** Regional age-related slopes extracted from linear mixed-effects models, accounting for subject- and node-level random effects. Panel **(a)** displays the distribution of age slopes for the cross-layer *Variation of Information (VI)*. Panel **(b)** displays the corresponding age slopes for the *Jaccard Distance*. **(c)** Three-dimensional glass brain rendering illustrating the spatial gradients of the Jaccard Distance age slopes. Subcortical hubs within striato-thalamo-cortical loops (putamen, pallidum, caudate, and thalamus) demonstrate the steepest positive age slopes (*p*_FDR_ *<* 0.05), whereas the limbic core (hippocampus and entorhinal cortex) exhibits flat to negative slopes.

### Topological Mechanism: Structural Constriction and Functional Expansion

To understand the geometric nature of this localized decoupling, we examined the lifespan trajectories of community degree—the total number of nodes co-assigned to a subcortical hub’s respective module. This revealed a striking multi-layer dissociation (Fig. 4 and Table 1).

**Table 1:**
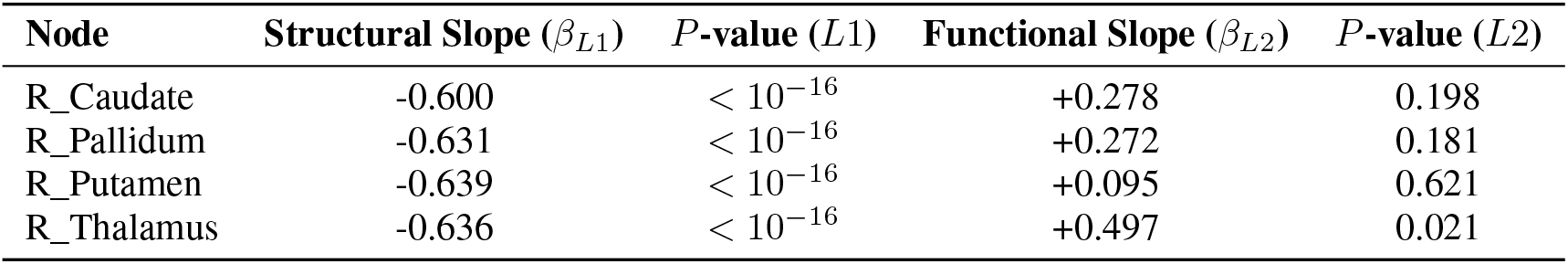
Lifespan Trajectories of Subcortical Community Degree.

**Figure 4:**
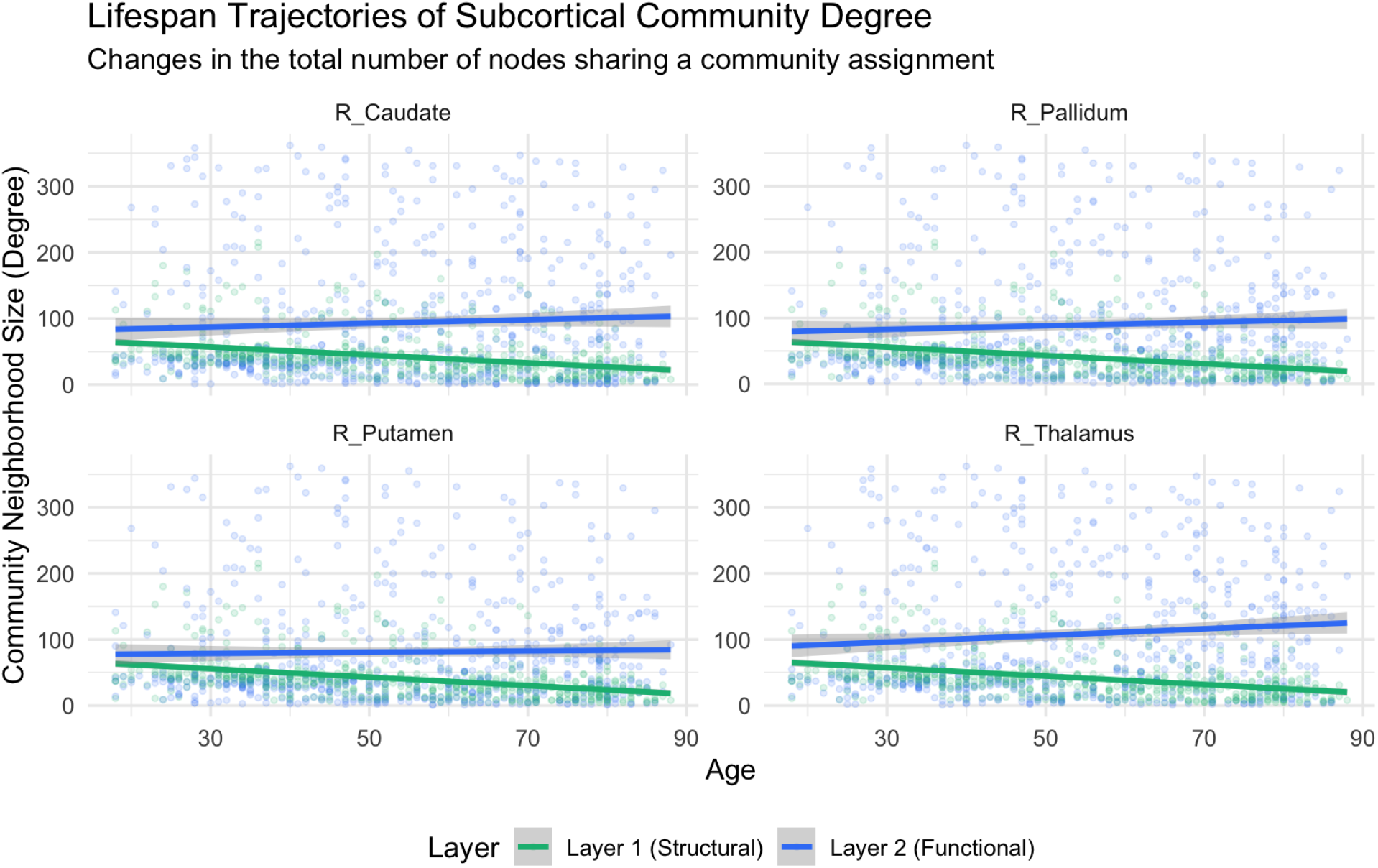
Lifespan Trajectories of Subcortical Community Degree. Structural neighborhoods (green) show highly significant linear constriction across all subcortical hubs with age, reflecting physical network isolation. In contrast, functional neighborhoods (blue) exhibit expanding or smearing boundaries, illustrating the geometric mismatch driving multi-layer divergence.

In the structural layer, all four subcortical epicenters exhibited severe and highly synchronized network constriction. Structural community degree decreased rapidly across all regions (e.g., Thalamus *β* = − 0.636, *p <* 10^−16^). This uniform collapse indicates a progressive physical isolation of subcortical nodes, likely driven by age-related white matter degradation and axonal pruning.

Conversely, the functional layer resisted this constriction, displaying uniformly positive age-related degree slopes. The thalamus demonstrated a statistically significant expansion in its functional community size (*β* = +0.497, *p* = 0.0217). This geometric mismatch—where functional boundaries expand and blur across a rapidly shrinking structural scaffold—mechanistically drives the progressive increase in Variation of Information.

### Pathway-Specific Reorganization: The Prefrontal-to-Limbic Shift

While global degree metrics capture network size, we performed an edge-wise tracking regression to identify the specific anatomical pathways driving this reorganization. Both layers exhibited a synchronized, highly specific topological shift, albeit at fundamentally different velocities.

First, we observed a profound frontostriatal and frontothalamic breakdown. With increasing age, all four subcortical hubs exhibited an accelerated reduction in co-assignment probability with higher-order cognitive control regions within the right dorsolateral prefrontal cortex (dlPFC), most notably areas R_9-46d, R_46, and R_8C (all *Q <* 10^−15^).

Simultaneously, this executive decoupling was offset by a “limbic hyper-consolidation.” The subcortical hubs demonstrated linearly increasing co-assignment with the primary allocortical core, including the piriform cortex (R_Pir), anterior cingulate (R_a24), amygdala, and entorhinal cortex (L_EC). Crucially, the velocity of functional reorganization outpaced structural adaptation. For example, the functional consolidation of the right putamen with the piriform cortex proceeded at more than double the velocity of its structural counterpart (*β* _*L*2_ = +0.0106 vs. *β* _*L*1_ = +0.0041).

### Global Divergence Acts as a Collinear Proxy for Cognitive Aging

Initial mass-univariate analyses revealed that global network divergence (Relative Information Loss) strongly correlated with both lower fluid intelligence (simple *r* = − 0.410, *p <* 0.001) (Figure 5) and slower reaction times (simple *r* = 0.282, *p <* 0.001). Crucially, no other cognitive domains in the broad Cam-CAN battery showed significant associations, providing strong discriminant validity that multiplex untethering specifically targets fluid problem-solving and motor adaptation rather than causing a ubiquitous cognitive shutdown.

**Figure 5:**
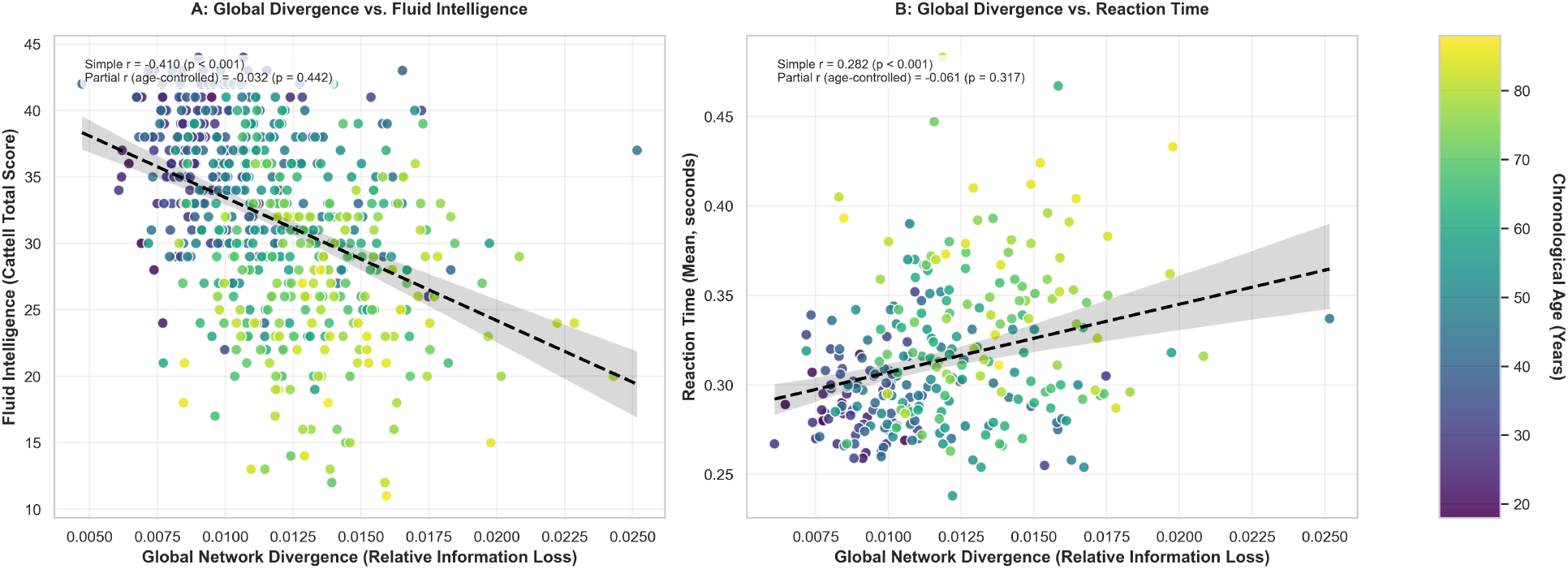
Global Divergence and Cognitive Performance. Scatter plots displaying the relationship between global network divergence (Relative Information Loss) and cognitive metrics, with chronological age represented by the color gradient. **(a)** Global divergence plotted against fluid intelligence (Cattell Total Score). The variables show a simple correlation of *r* = − 0.410, which attenuates to a partial correlation of *r* = − 0.030 after controlling for age. **(b)** Global divergence plotted against reaction time. The variables show a simple correlation of *r* = 0.282, which attenuates to a partial correlation of *r* = −0.047 after controlling for age.

However, upon implementing partial correlations to control for chronological age, these global associations largely attenuated (Table 2). For example, the relationship between relative information loss and fluid intelligence (Cattell Total Score) was no longer significant after controlling for age (*r*_*partial*_ = − 0.030, *p* = 0.508), and reaction time showed a similarly attenuated association (*r*_*partial*_ = − 0.047, *p* = 0.437). This indicates that, at the whole-brain level, structural-functional untethering operates collinearly with chronological aging, acting as a generalized biological marker for global cognitive decline rather than an independent driver.

**Table 2:** Global Network Divergence: Age-Controlled Partial Correlations.

| Domain | Partial Correlation (controlling for age) |  |
| --- | --- | --- |
|  | rel_loss | JSD |
| **Fluid Intelligence (Cattell)** |  |  |
| Total Score | $r = -0.030$ ($p = 0.508$) | $r = 0.027$ ($p = 0.508$) |
| SubScore 1 | $r = 0.005$ ($p = 0.896$) | $r = 0.045$ ($p = 0.280$) |
| SubScore 2 | $r = -0.028$ ($p = 0.501$) | $r = -0.004$ ($p = 0.918$) |
| SubScore 3 | $r = -0.004$ ($p = 0.929$) | $r = 0.030$ ($p = 0.464$) |
| SubScore 4 | $r = -0.067$ ($p = 0.103$) | $r = 0.010$ ($p = 0.804$) |
| **Motor Learning** |  |  |
| Late Post-Exposure Trajectory Error Mean | $r = -0.106$ ($p = 0.081$) | $r = -0.098$ ($p = 0.105$) |
| Early Post-Exposure Movement Time Mean | $r = 0.068$ ($p = 0.263$) | $r = 0.069$ ($p = 0.258$) |
| Reaction Time Mean | $r = -0.061$ ($p = 0.317$) | $r = -0.047$ ($p = 0.437$) |
| Late Exposure Trajectory Error SD | $r = 0.067$ ($p = 0.268$) | $r = 0.119$ ($p = 0.049$) |
| *Note:* Partial correlations controlling for chronological age ($N = 589$). None of the associations survive False Discovery Rate (FDR) correction for multiple comparisons ($q > 0.05$). This demonstrates that global network divergence operates collinearly with chronological age, acting as a proxy for general aging rather than an independent driver of cognitive decline. | | |

## 3 Discussion

Our study establishes a new organizing principle of healthy aging: a progressive information divergence between the brain’s structural scaffold and its functional traffic. By leveraging a multiplex network framework on the Cam-CAN dataset, we demonstrate that the aging process is not merely defined by the degradation of isolated components, but by a systematic “untethering” of functional dynamics from their underlying white-matter constraints. This global decoupling—most pronounced in subcortical “switchboards” yet resisted by the limbic core—offers a nuanced view of how the human brain maintains, and eventually loses, its communicative efficiency across the adult lifespan.

### The Untethering of Structure and Function

The central hallmark of the aging connectome identified here is the linear increase in Jensen-Shannon Divergence (*d*_JS_) and relative information loss (*q*(*C*)) with age. Our findings significantly advance the multi-modal connectomics landscape, which has historically relied on static or single-layer paradigms. Foundational multiplex frameworks have previously demonstrated that white-matter anatomy non-trivially constrains functional configurations via layer edge-motifs Battiston et al. [2017], macroscale hubs Battiston et al. [2018], and distinct layer-specific assortativity profiles Lim et al. [2019]. Furthermore, recent developments confirm that structure-function relationships are deeply embedded within community-level architectures Puxeddu et al. [2022] and indirect structural path bandwidths Parsons et al. [2022]. However, these frameworks have predominantly evaluated young, static cohorts. Shifting this paradigm to a lifespan perspective, recent aging literature has begun utilizing cross-modality frameworks to track age-related declines in structural-functional similarity Jauny et al. [2024], macroscale core migrations Devrome et al. [2023], and altered integrated rich-club hierarchies Khalilian et al. [2024].

While these recent aging studies successfully capture a general decline in cross-modality alignment, they remain methodologically bounded by first-order, strictly linear edge-to-edge Pearson correlations. Such linear coupling approaches are blind to higher-order topological geometry, complex non-linear network distributions, or the shared entropy between layers. By introducing distribution-free information-theoretic metrics based on graph Laplacian density matrices, our framework quantifies the true statistical distance and irreducibility of the integrated connectome.

In younger cohorts, the functional connectome (FC) remains closely aligned with the structural connectome (SC), suggesting that neural communication is highly efficient and strictly guided by physical wiring. As the brain matures and ages, this alignment erodes. The progressive increase in divergence indicates that functional interactions increasingly traverse topological pathways that are not strictly governed by the underlying deterministic structural fiber bundles. This shift likely reflects a transition from a structurally constrained mode of communication to a more degenerate functional state. While this “untethering” represents a significant departure from the youthful connectome, it also necessitates a shift in how we model the aging brain. Our findings show that by the eighth decade, the information loss associated with collapsing these layers into a single aggregate nearly doubles. This demonstrates that a multiplex architecture is not merely a mathematical convenience but a biological necessity; a single-layer view fails to capture the growing independence of functional traffic that characterizes the aging process.

Our findings also speak directly to developmental accounts of SC-FC coupling, which have shown that structure-function alignment strengthens across childhood and adolescence in parallel with gains in intelligence and executive function (Soman et al. [2023], Baum et al. [2020], Feng et al. [2024]). Rather than continuing this trajectory, our data indicate that healthy aging is characterized by a reversal: the coupling established over development progressively erodes across adulthood, suggesting that SC-FC coupling may follow an inverted-U trajectory across the human lifespan — rising through youth, and declining thereafter. Critically, however, this reversal is not uniform across the brain. While decoupling in subcortical switchboards parallels the decline of fluid intelligence and motor adaptation, the limbic core resists this reversal, actively preserving the same close structure-function alignment associated with cognitive competence earlier in life. This regional dissociation refines the developmental account: it is not a single, global coupling-cognition relationship that determines cognitive outcomes in aging, but rather which specific circuits retain versus relinquish structure-function fidelity.

### Functional Specificity and Discriminant Validity

A central question regarding age-related brain network degradation is whether it causes a uniform deterioration of cognition or a highly selective vulnerability. Our behavioral results strongly support the latter. Out of a comprehensive battery of cognitive and behavioral assessments screened across the Cam-CAN repository, significant raw associations with our network metrics were strictly localized to two specific domains: Fluid Intelligence (Cattell’s Culture Fair total score) and Motor Learning (Visuomotor Rotation task performance).

This behavioral specificity aligns perfectly with our anatomical findings; both affected domains rely heavily on high-fidelity signal routing through the striato-thalamo-cortical scaffold—the exact regions we identified as the primary epicenters of multiplex untethering. Sparing cognitive domains like crystallized knowledge, this striking discriminant validity confirms that structural-functional information divergence is not merely diffuse biological noise, but a targeted topological breakdown of the neural loops necessary for fluid cognitive adaptation and real-time behavioral flexibility.

### Subcortical “Switchboards” as the Epicenter of Aging

Our node-level analysis identifies subcortical hubs—specifically the putamen, pallidum, caudate, and thalamus—as the primary drivers of age-related divergence. These regions serve as the brain’s central switchboards, integrating sensory-motor and cognitive information from across the cortex. In the context of established neurocognitive aging literature, this untethering aligns strongly with the frontostriatal aging hypothesis. The basal ganglia are notably susceptible to age-related dopaminergic degeneration, characterized by a marked decline in striatal dopamine transporters van Dyck et al. [2002] and widespread alterations in midbrain dopaminergic regulation Dreher et al. [2008]. We propose that the steep structural-functional decoupling observed here provides a macroscopic connectomic signature for this localized neurochemical vulnerability. This topological “rerouting” offers an empirical network-level explanation for the degradation of processing speed and fluid problem-solving in cognitive aging Salthouse [1996].

Strikingly, this subcortical vulnerability stands in direct contrast to the remarkable multiplex stability of the limbic core, particularly the hippocampus and entorhinal cortex. Prior lifespan studies employing conventional edge-wise correlations reported an complete absence of age-related alterations within the connection strengths of the limbic system Khalilian et al. [2024]. By moving past first-order correlations and tracking fine-grained regional community partitions via the multiplex map equation, we show that this flat trajectory is not an absence of signal, but rather a profound biological phenomenon driven by distinct multi-layer geometric constraints.

### The Geometry of Untethering: Shrinking Scaffold vs. Expanding Traffic

An apparent paradox emerges from our edge-wise and neighborhood analyses: how can the structural and functional layers drift apart globally (increasing Variation of Information) while simultaneously executing a shared directional migration—namely, a systematic decoupling from the prefrontal cortex and a hyper-consolidation within the limbic core? Our microscale and mesoscale findings resolve this paradox through two distinct mechanisms: a profound layer-specific velocity mismatch and a severe geometric dissociation in community neighborhood size (Degree).

First, our edge-wise tracking reveals that while both layers are withdrawing from the executive prefrontal cortex (e.g., area `R_9p`) and retreating into the primitive allocortical core (e.g., `R_Pir`), they do so at radically out-of-sync velocities. Subcortical structural connections to the dorsolateral prefrontal cortex degrade at a significantly faster rate (*β* _*L*1_ = −0.0151) than functional traffic withdraws (*β* _*L*2_ = −0.0107). Conversely, the functional hyper-consolidation of these hubs into the limbic system occurs at more than double the rate of physical structural remodeling (*β* _*L*2_ = +0.0106 versus *β* _*L*1_ = +0.0041). Because the structural substrate and functional signaling operate on fundamentally mismatched temporal trajectories, they inevitably drift into global topological misalignment (*d*_JS_).

Second, this untethering is driven by a fundamental geometric crossover in community size, as visualized in (Fig. 4). Structurally, subcortical epicenters experience severe, highly synchronized network constriction ((*β ≈* − 0.63, *p <* 10^−16^), indicating a progressive physical isolation driven by white-matter axonal pruning. Functionally, however, these hubs completely resist this constriction. Instead, their functional communities display positive age-slopes, with the right thalamus demonstrating a statistically significant neighborhood expansion (*β* = +0.497, *p* = 0.0217).

This pattern directly captures a state of functional de-differentiation and a loss of neural segregation. While the structural scaffold shrinks and isolates the subcortex, the functional layer expands its boundaries, engaging in unconstrained cross-talk to compensate for the failing physical skeleton. This structural-functional crossover confirms that the observed divergence is not an artifact of passive tissue decay or random atrophy, but a highly coordinated, multi-layer adaptive phenomenon.

### The Resilient Limbic Core: Fidelity vs. Flexibility

In sharp contrast to the structural constriction and functional smearing observed in subcortical hubs, the preserved structural-functional coupling in the limbic system can be interpreted through the lens of the Scaffolding Theory of Aging and Cognition (STAC) Park and Reuter-Lorenz [2009], which posits that the brain engages continuous compensatory neural mechanisms to protect cognitive function against structural degradation. We propose that the active maintenance of the structural-functional tether in the limbic system represents a fundamental biological scaffold.

While subcortical regions exhibit progressive untethering, the limbic core prioritizes “representational fidelity” over “computational flexibility.” Despite age-related declines in episodic memory performance, these memory-critical regions actively maintain high structural-functional community alignment across the lifespan. This unique resilience suggests a fundamental biological imperative: while subcortical circuits may tolerate or even benefit from functional reorganization, the limbic system prioritizes strict topological persistence to protect its core representational architecture from catastrophic information corruption. In this light, the preserved limbic anchor acts as a biological barrier against the chaotic neural disorganization characteristic of Alzheimer’s disease. As these regions are established epicenters for early tau pathology, the eventual breakdown of this specific multiplex anchor may serve as a potent connectomic biomarker for the transition from normative aging to neurodegeneration.

### Statistical Robustness and Methodological Considerations

Methodologically, our work underscores the necessity of multiplexity in connectomics. By using the Multiplex Map Equation for flow-based community detection, we tracked how functional “traffic” persists or exits across structural layers—a level of meso-scale detail invisible to single-modality approaches.

It is vital to contextualize the statistical magnitude of the observed global divergence and its connection to human performance. In our cross-sectional cohort of 589 healthy adults, chronological age explained a substantial portion of the variance for both relative information loss (*R*^2^ = 0.36) and Jensen-Shannon divergence (*R*^2^ = 0.17). On a global scale, mass-univariate Pearson correlations initially linked this information loss to lower fluid intelligence (*r* = − 0.410) and slower reaction times (*r* = 0.282). However, these global brain-behavior associations attenuated when controlling for chronological age (partial correlations: *r* = − 0.030 and *r* = − 0.047, respectively). This demonstrates that at the whole-brain level, global network untethering operates collinearly with aging, acting as a generalized biological marker for global cognitive slowing rather than an independent driver.

### Limitations and Future Directions

Several limitations warrant consideration. First, as a cross-sectional study, we cannot definitively rule out cohort effects; future longitudinal multiplex studies will be essential to confirm individual trajectories of structural-functional decoupling.

Second, and critically, our functional connectivity estimates are derived from resting-state fMRI, which captures intrinsic, task-independent fluctuations. While resting-state paradigms offer excellent standardization and comparability across the lifespan, they may underestimate the full extent of structure-function coupling that manifests during active task engagement. Motor learning and fluid intelligence tasks inherently require the recruitment of task-specific functional networks that may be more tightly constrained by structural wiring than the resting state.

Future work should therefore implement task-based fMRI paradigms—particularly visuomotor adaptation tasks and fluid reasoning paradigms—concurrently with structural connectomics. Such designs could reveal stronger structure-function associations and may unmask task-specific decoupling patterns that remain latent during rest. This is particularly relevant for the subcortical epicenters identified here: the putamen, pallidum, caudate, and thalamus are most engaged during active motor and cognitive processing, not at rest. Task-based multiplex analyses may therefore provide more sensitive biomarkers of age-related functional reorganization.

Third, while we have successfully mapped the spatial topography of age-related structural-functional divergence, the causal mechanisms underlying this untethering remain speculative. The link to dopaminergic degeneration, while plausible, requires direct validation through multimodal imaging (e.g., PET with dopaminergic tracers) or pharmacological manipulation.

Finally, while this study establishes a baseline across the healthy adult lifespan, a vital future frontier lies in clinical translation. Recent multi-layer brain network applications in clinical connectomics have successfully pinpointed altered cross-layer participation in restless legs syndrome Park et al. [2024], disrupted indirect white-matter routing bandwidths in schizophrenia Li et al. [2024], and sensory-modality uncoupling in preterm neonates Quinones et al. [2025]. Testing whether our specific subcortical divergence patterns and information-theoretic indices can distinguish normative human aging from pathological clinical neurodegeneration is a critical next step—particularly in prodromal Parkinson’s disease or Alzheimer’s disease, where subcortical and limbic circuits respectively serve as early pathological targets.

### Conclusion

Healthy aging is characterized by a progressive structural-functional drift that profoundly reconfigures the brain’s information architecture. By identifying subcortical hubs—putamen, pallidum, caudate, and thalamus—as the focal points of this drift and the limbic system as its stable anchor, we provide a new framework for understanding neural resilience. These findings reveal that the hallmark of the aging brain is not merely the loss of connectivity, but a fundamental shift in the governing relationship between the physical and functional layers of the human connectome. By establishing this normative multiplex baseline, our work provides a foundation for distinguishing healthy cognitive aging from the early topological signals of neurodegenerative disease, with the important caveat that future task-based and longitudinal studies will be essential to validate and extend these findings.

## 4 Materials and Methods

### 4.1 Data Acquisition and Cohort Characteristics

We utilized multimodal neuroimaging data from the Cambridge Centre for Ageing and Neuroscience (Cam-CAN)Shafto et al. [2014], Taylor et al. [2017]. The final cohort consisted of *N* = 589 healthy adults (302 female, 288 male) spanning the adult lifespan (ages 18–88). Imaging was performed on a 3T Siemens TIM Trio system, including T1-weighted and T2-weighted structural MRI, diffusion-weighted imaging (DWI), and resting-state functional MRI (rs-fMRI). Detailed acquisition parameters and exclusion criteria are provided in the Supplementary Information.

### 4.2 Connectome Construction

#### Structural Connectivity (SC)

Cortical and sub-cortical parcellation was performed using the HCP-MMP 1.0 atlas (379 regions)Glasser et al. [2016] applied via FreeSurfer Fischl [2012]. Structural connectomes were generated using MRtrix3, employing anatomically constrained tractography (ACT) with 10 million streamlinesTournier et al. [2019]. Edge weights were refined using the SIFT2 algorithm to ensure biological accuracy of white-matter fiber densities.

#### Functional Connectivity (FC)

Functional data were preprocessed for motion correction, denoising, and registration. For each of the 379 regions of interest (ROIs), representative time series were extracted by computing the first principal component (principal eigenvariate) of the signals across all voxels within the respective regional mask. Pairwise functional connectivity was subsequently estimated using the mutual information of these blood-oxygen-level-dependent (BOLD) signals.

### 4.3 Information-Theoretic Global Measures in Multiplex

We adopt an information-theoretic perspective on the SC-FC multiplex network. A key advantage of viewing the brain as an information-theoretic module is that it shifts the analysis of multiplex networks from purely topological descriptions (e.g., comparing edge counts or clustering coefficients) to a principled, scale-invariant framework that quantifies how information is distributed, shared, and compressed across layers. In the context of our two-layer multiplex network—structural connectivity (SC) and functional connectivity (FC)—this perspective offers three specific benefits: it captures statistical differences between layers without assuming linearity or specific distributions; (ii) it determines whether the two layers carry redundant or complementary information; and (iii) it identifies community structures that optimally explain information flow across both layers. These benefits directly motivate our choice of measures. We therefore analyze the SC-FC multiplex using Jensen–Shannon divergence to quantify the distinguishability between layers, multiplex reducibility to assess whether the multiplex can be collapsed into a single layer without information loss, and the Multiplex Map Equation algorithm (Infomap) to detect communities that integrate structure and function.

We represent the brain as a two-layer multiplex network *M* ={*L*_SC,_ *L*_FC_}, where nodes correspond to identical anatomical regions across both layersKivelä et al. [2014], De Domenico et al. [2013], Boccaletti et al. [2014]. The system is represented as a supra-adjacency matrix 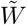, capturing the intra-layer connectivity across the diagonal blocks and the the inter-layer coupling between structural scaffolds and functional traffic along the off-diagonal blocks Boccaletti et al. [2014]. The off-diagonal blocks are identity matrices, to represent the same set of brain regions interacting across both layers.

To quantify the age-related changes of these layers, we primarily adopt two information-theoretic metrics:

1. **Jensen-Shannon Divergence (***d*_**JS**_): From an information standpoint, we first ask: how different are the connectivity distributions of structural connectivity (SC) and functional connectivity (FC)? To answer this, we employ the Jensen–Shannon divergence, a symmetric information-theoretic measure that quantifies the statistical distance between the two layers without assuming linearity or any particular distribution. A larger Jensen–Shannon divergence indicates that SC and FC carry fundamentally different information; conversely, a smaller divergence suggests that the two layers are statistically similar and share most of their information. The Jensen–Shannon divergence requires inputs to be positive semidefinite with unit trace (density matrices), ensuring all eigenvalues are non-negative and sum to one—analogous to a probability distribution. The graph Laplacian is inherently positive semidefinite; after rescaling to unit trace, it serves as a valid density matrix that encodes the connectivity topology of each layer. Hence, we construct rescaled Laplacian density matrices (*ρ, σ*) for the SC and FC layers. The J-S divergence is defined as:

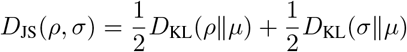

where 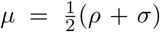 is a mixture defined as in De Domenico et al. [2015b]. The resulting distance 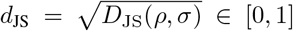 quantifies the dissimilarity between the structural topology and functional dynamics De Domenico et al. [2015b].
2. **Multiplex Reducibility (*q*(*C*)):** A central question in multiplex network analysis is whether the distinct layers—here, structural connectivity (SC) and functional connectivity (FC)—carry complementary information or whether they can be merged into a single aggregated network without significant loss. To address this, we employ the concept of multiplex reducibility, which quantifies the information-theoretic distance between the full multiplex and its aggregated counterpart. This distance is grounded in the Von Neumann entropy (De Domenico et al. [2015b]) of each layer’s rescaled Laplacian density matrix. Just as Shannon entropy measures the uncertainty of a probability distribution, the Von Neumann entropy *S*(*ρ*) = − Tr(*ρ* ln *ρ*) measures the spectral entropy of a network, capturing how evenly information is distributed across its eigenmodes. By comparing the Von Neumann entropy of the multiplex (treating layers as independent) with that of the aggregated network (layers collapsed), we determine whether merging the layers results in information loss. A low reducibility (i.e., a large information-theoretic distance) indicates that SC and FC are statistically distinct and cannot be reduced to a single layer without discarding meaningful information—justifying their treatment as a genuine multiplex. Using the Von Neumann entropy, we evaluate the relative information loss incurred by collapsing the multiplex into a single-layer aggregate network (De Domenico et al. [2015b]). The relative entropy *q*(*C*) is defined as:

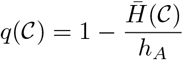

where 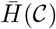 is the average entropy of the multiplex layers and *h*_*A*_ is the entropy of the aggregated graph obtained by shrinking the multiplex to a single network. A higher *q*(*C*) indicates that the multiplex structure carries significantly more information than the aggregate network De Domenico et al. [2015b].

### 4.4 Meso-scale Dynamics

Beyond measuring layer distinguishability, reducibility, and edge overlap, we seek to understand the meso-scale organization of the SC-FC multiplex—specifically, how brain regions cluster into functional modules when considering structure and function jointly. We identified communities using the multiplex extension of the map equation De Domenico et al. [2015c], which defines communities as groups of state nodes that trap random-walk flow within and across layers for relatively long times. In this framework, each physical node has a layer-specific state node, and community detection is performed by finding the partition that minimizes the description length of the random walk on the multiplex. When empirical interlayer links are unavailable, cross-layer movement is controlled by a relax rate *r*: with probability 1 − *r*, the walker follows intralayer edges, and with probability *r*, it may switch layers through the corresponding physical node. We estimated *r* from simulations designed to reflect our data structure, in which layers shared partially aligned but nonidentical community organization under sparse connectivity. Community recovery was best for *r ∈* [0.15, 0.3]; therefore, we used *r* = 0.15 in the main analysis. An in-depth discussion is provided in the Supplementary Information.

Node-level reorganization was quantified by comparing the community assignment vectors of each region across the SC and FC layers using:

- **Variation of Information (VI):** An information-theoretic distance measuring the uncertainty remaining in one partition after observing the other Meil ă [2007].
- **Jaccard Distance:** A metric measuring the dissimilarity between node-specific community memberships Jaccard [1901], Levandowsky and Winter [1971].

For each subject and node, we quantified structure–function community mismatch using the normalized variation of information between structural and functional community allocation profiles. Age-related effects were assessed using a linear mixed-effects model with age as a fixed effect and subject- and node-level random effects to account for repeated measurements and node-specific variability. To localize age-related changes, we also performed node-wise linear regressions of this measure on age and controlled for multiple testing using the false discovery rate.

#### Geometrical and Pathway-Specific Reorganization

To evaluate the geometrical nature of network reorganization, we computed the *community degree* for each target subcortical node, defined as the total number of anatomical regions co-assigned to its layer-specific community. Lifespan trajectories of structural and functional community degree were modeled using linear regression to distinguish network constriction from boundary expansion.

Furthermore, to identify the precise anatomical pathways driving this reorganization, we performed an edge-wise tracking regression. For each subcortical hub, we quantified the binary probability of community co-assignment with all other 378 cortical and subcortical regions across the lifespan. Node-to-node co-assignment probabilities were regressed against age for both the SC and FC layers independently, followed by a False Discovery Rate (FDR) correction (*q <* 0.05) to isolate statistically significant functional decoupling (negative slopes) and limbic recruitment (positive slopes).

### 4.5 Behavioral Data and Discriminant Validity

To investigate the functional consequences of multiplex information divergence, we analyzed behavioral data from the Cam-CAN repository. To establish discriminant validity and ensure our divergence metrics were not merely capturing a non-specific global decline, we initially screened a comprehensive battery of cognitive and behavioral assessments. Significant raw associations with our network metrics were strictly localized to two domains: Fluid Intelligence (measured via Cattell’s Culture Fair test) and Motor Learning (measured via the Visuomotor Rotation task). Given that both domains heavily rely on striato-thalamo-cortical loops—the precise anatomical epicenters of multiplex untethering identified in our model—these two measures were selected for in-depth regional analysis.

#### 4.5.1 Hierarchical Statistical Pipeline

We employed a hierarchical statistical approach. First, mass-univariate Pearson correlations were used to identify global baseline associations. Next, partial correlations controlling for chronological age were implemented to distinguish age-independent functional relationships from generalized chronological decline.

## Author Contributions

DR, MD and DG conceived and designed the project. DG, DR, and MD performed the data analysis. MD provided overall supervision for the study. DR and DG drafted the initial manuscript. MD and LU contributed to the critical revision and editing of the manuscript. All authors have read and approved the final version of the manuscript.

## Data and Code Availability

The complete scripts used for data preprocessing, and network analysis are openly available in the GitHub repository at https://github.com/dipanjan-neuroscience/CamCan_Str_Func.

## Acknowledgement

Data used in this work were obtained from the CamCAN repository (https://cam-can.mrc-cbu.cam.ac.uk/), based on the CamCAN study [Shafto et al., 2014].

## Supplementary Information

### S1 Detailed Neuroimaging Acquisition Parameters

All data were acquired at the Medical Research Council Cognition and Brain Sciences Unit on a 3T Siemens TIM Trio System.

- **T1-weighted (T1w) MRI:** 3D MPRAGE sequence; TR = 2250 ms, TE = 2.99 ms, TI = 900 ms, flip angle = 9^°^; FOV = 256 *×* 240 *×* 192 mm; 1 mm isotropic voxels; GRAPPA acceleration factor = 2.
- **T2-weighted (T2w) MRI:** 3D SPACE sequence; TR = 2800 ms, TE = 408 ms, TI = 900 ms; FOV = 256 *×* 256 *×* 192 mm; 1 mm isotropic voxels; GRAPPA = 2. **Diffusion-Weighted Imaging (DWI):** 2D twice-refocused SE-EPI sequence; TR = 9100 ms, TE = 104 ms; FOV = 192 *×* 192 mm; 66 axial slices (2 mm isotropic); multi-shell b-values = 0, 1000, 2000 s/mm^2^ across 30 directions. Total readout time = 0.0684 s (echo spacing = 0.72 ms).
- **Resting-state fMRI (rs-fMRI):** T2^*∗*^-weighted GE-EPI; TR = 1970 ms, TE = 30 ms, flip angle = 78^°^; 261 volumes; 32 axial slices; voxel size = 3 *×* 3 *×* 4.44 mm; FOV = 192 *×* 192 mm.

### S2 Structural and Functional Preprocessing Pipelines

#### S2.1 Functional Connectivity Pipeline

##### Image Preprocessing

Resting-state fMRI (rs-fMRI) data preprocessing was performed using SPM12 (Statistical Parametric Mapping, Wellcome Centre for Human Neuroimaging, London, UK). To ensure magnetization stabilization, the first 5 volumes of each functional session were discarded as dummy scans. The remaining 256 volumes underwent slice-timing correction to account for interleaved acquisition delays, utilizing a sequential descending slice order (32 slices, TR = 1.97 s, TA = 1.9084 s) with the first slice selected as the reference.

Following slice-timing correction, the functional volumes were realigned to their mean image using a rigid-body transformation to estimate and correct for head motion. The high-resolution T1-weighted (T1w) structural image was then co-registered to the mean functional image using a Normalized Mutual Information (NMI) cost function. Unified segmentation was applied to the co-registered structural scan to separate brain tissues into gray matter, white matter (WM), and cerebrospinal fluid (CSF) using SPM12’s tissue probability maps. The resulting forward deformation fields were used to spatially normalize the functional images into the Montreal Neurological Institute (MNI) standard space, resampled to 2 *×* 2 *×* 2 mm isotropic voxels. Finally, spatial smoothing was executed using an anisotropic Gaussian kernel with a Full Width at Half Maximum (FWHM) of 6 *×* 6 *×* 8 mm.

To minimize confounding impacts from head motion and physiological fluctuations, a two-step General Linear Model (GLM) framework was deployed. A set of 18 motion-related regressors was constructed, comprising the 6 rigid-body realignment parameters, their immediate preceding (1-back) temporal derivatives, and their squared values. In addition, physiological noise characteristics were captured by extracting the mean BOLD time series from 6 mm radius spherical Volumes of Interest (VOIs) centered within the white matter (MNI coordinates: [0, −24, −33]) and cerebrospinal fluid (MNI coordinates: [0, −40, −5]), bounded by the subject-specific whole-brain mask.

These 20 nuisance variables (18 motion parameters + 1 WM time series + 1 CSF time series) were included as multiple regressors in a secondary GLM estimation configuration. Concurrently, a high-pass filter with a cutoff period of 128 s was incorporated into the design matrix to systematically remove low-frequency scanner drifts. The residuals from this regression model were retained as the denoised BOLD signal for network evaluation.

##### Network Construction

Following the pre-processing steps, the denoised BOLD signals were used for network construction. Specifically, the brain was subdivided into regions of interest using a whole-brain parcellation namely the HCP-MMP 1.0 atlas Glasser et al. [2016] and the first principal component of the voxel-wise BOLD signals was extracted for each parcel. These regions became the nodes in the network while the edges among them were computed using the mutual information between each pair of the time series, to capture both the linear and non-linear dependencies. These pairwise values were organized into a weighted, symmetric adjacency matrix representing the subject-specific functional connectivity matrix.

#### S2.2 Structural Connectivity Pipeline

##### Image Preprocessing

For each subject, we first used the T1-weighted (T1w) anatomical MRI to create the parcellation (a set of brain regions) that would later serve as the nodes in the structural connectomes. This was implemented using FreeSurfer Fischl [2012], which takes a T1w scan and reconstructs the cortical surfaces and segments the sub-cortical structures and produces the whole-brain anatomical segmentation. The cortical and sub-cortical regions were defined using the HCP-MMP 1.0 atlas Glasser et al. [2016] by transferring the atlas labels onto each subject’s anatomy. The resulting parcellation was used with MRtrix3 Tournier et al. [2019] for subsequent connectome generation.

Subsequently, for each subject, we used the diffusion MRI (DWI) scan to estimate the main directions of white-matter pathways and to generate tractography. This part of the analysis was run using MRtrix3 Tournier et al. [2019], with some standard correction steps, such as correction for acquisition issues (such as noise and head movement) that are carried out using FSL and ANTs. Additionally, a brain mask was created so that the subsequent processing was restricted to the brain tissue. Next, using MRtrix3, we estimated the directions of the fiber tracts from the corrected diffusion data and prepared the images required for tractography. Here, we also used the T1w scan as an anatomical reference for the tractography by aligning it to the diffusion scan and the resulting tissue map was used by MRtrix3 during the construction of the tractography.

Finally, whole-brain tractography was performed in MRtrix3 to generate a tractogram containing 10 million streamlines, and the streamline set was refined to obtain the weights using MRtrix3 SIFT2 for each streamline.

##### Network Construction

To construct the structural connectomes for each of the subjects, we took the parcellations from the whole brain to be the set of nodes and derived the edges using the set of reconstructed white matter pathways from the tractogram. Each element of the matrix represents the strength of the connection, based on the number of the white matter pathways (i.e. streamlines starting in one parcel and in ending in another) between each pair of brain regions. This yielded a symmetric, undirected connectivity matrix for each subject in the study.

We retained the strongest 10% edges in both the structural and functional connectivity matrices to minimize the impact of weak and noisy connections in our analyses.

### S3 Multiplex Network Analysis

A closer look at most natural and social phenomena shows that entities rarely act in isolation; instead, they share intricate webs of interconnections. A common approach for representing these interconnections in the network is the use of graphs where these relationships are typically represented such that the entities are treated as nodes or vertices and the links between them as edges Newman [2018]. More recently, multilayer network analysis (Kivelä et al. [2014],)

De Domenico et al. [2013] has emerged as a powerful framework for capturing the added complexity of such systems, enabling a multi-view lens on the underlying mechanisms. By modeling multiple types of interactions among the same physical entities across layers, such as at different time points or across modalities, and more generally allowing the layers to represent interconnected node sets, this framework offers deeper insights into how interdependent components interact across contexts within a system.

A *Multilayer Network* as defined in Boccaletti et al. [2014] can be represented as a pair 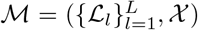, where ℒ _*l*_ = (*U*_*l*_, *F*_*l*_) denotes the *l*-th layer with node set 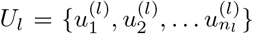 and intra-layer edge set *F*_*l*_ *⊆ U*_*l*_ *× U*_*l*_. The layers may be directed or undirected, and may carry weights. The inter-layer connections are collected in *X* = {*F*_*lm*_ *⊆ U*_*l*_ *× U*_*m*_ : *l, m* ∈ {1, 2, … *L*}, *l* ≠ *m*}.

For each layer, the within-layer adjacency matrix is 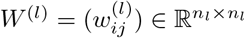, with

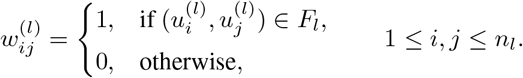

Likewise, the inter-layer adjacency between layers *l* and *m* is 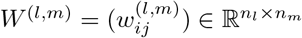, with

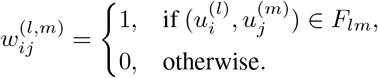

Stacking all intra- and inter-layer blocks yields the *supra-adjacency matrix*

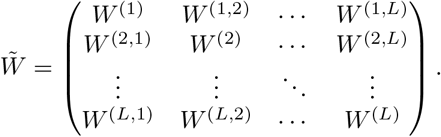

This block representation is also referred to as a flattened or matricized form of the multilayer system.

A *Multiplex Network* defined in Boccaletti et al. [2014] is a special case where node sets are identical across layers: *U*_1_ = *U*_2_ = · · · = *U*_*L*_ = *U*. Cross-layer edges connect only a node to its replicas in other layers, i.e., *F*_*lm*_ = {(*u, u*) : *u* ∈ *U*} for every *l* ≠ *m*. While the node set is common, intra-layer edge sets *F*_*l*_ may differ across layers.

#### S3.1 Multiplex Summary Measures

Here we explore certain information-theoretic *global* measures of the multiplex network, i.e., quantities that provide a system-level view by aggregating cross-layer complexity into “information”-based summaries. The brain is fundamentally an information processing system and the primary question in this study is to examine how the structural “map” and the functional “traffic” coordinate as a coupled system to support information processing across the healthy adult lifespan. Information-theoretic global measures are well suited to this aim because they provide entropy-based summaries of uncertainty, redundancy, and cross-layer divergence, thereby capturing structure–function integration using a single quantity.

##### Jensen-Shannon Distance

Let *D*^(*l*)^ and *W* ^(*l*)^ denote the degree matrix and adjacency matrix of layer *L*_*l*_, *l* ∈ {1, 2, …, *L*} respectively. One can then construct the Laplacian matrix as Λ^(*l*)^ = *D*^(*l*)^ − *W* ^(*l*)^. A rescaled Laplacian or a density matrix is obtained from the same by dividing the Laplacian by 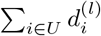, where 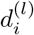 denotes the degree of node *i*. Given two density matrices *ρ* and *σ*, it can be shown that the Kullback-Leibler (KL) divergence is given by

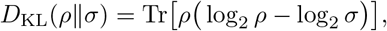

which is not a symmetric metric in general. The Jensen-Shannon distance between the graphs has been defined by De Domenico et al. [2015b], by first defining the mixture

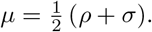

The Jensen-Shannon Distance is the average of the two KL divergences to the mixture

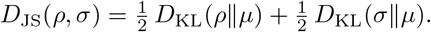

The Jensen-Shannon distance is reported as

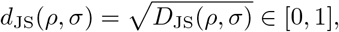

which is symmetric, and equals 0 if *ρ* = *σ*, and increases as the two matrices differ.

##### Reducibility in Multiplex Networks

The aim here is to quantify the distinctiveness of a multiplex representation from its aggregate representation (obtained by adding all the adjacency matrices and compressing the structure to a single layer network) using Von-Neumann entropy, which depends on the number of layers as well as the structure of those layers. In a multiplex network with *L* layers and the set of *N × N* adjacency matrices given by *M* = {*W* ^(1)^, *W* ^(2)^, …, *W* ^(*L*)^}, the Von-Neumann entropy *H*(*M*) of a multiplex network is defined as the sum of the Von-Neumann entropies of all its layers i.e. 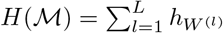, where 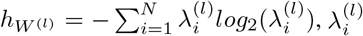, are the eigenvalues of the density matrix obtained by rescaling the Laplacian Λ^(*l*)^.

The general idea is that suppose from the multiplex network = *W* ^(1)^, *W* ^(2)^, …, *W* ^(*L*)^, one obtains a reduced multiplex structure = *C*^(1)^, *C*^(2)^, …, *C*^(*K*)^, with *K ≤ L* layers, where the adjacency matrix *C*^(*l*)^, *l ∈* 1, 2, …, *K*, is one that is obtained from one of the adjacency matrices of the original multiplex or is obtained by compressing two or more adjacency matrices from the same. Then, the entropy per layer of this multiplex *C* is given by

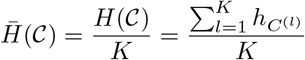

Let *h*_*A*_ denote the Von-Neumann entropy of the aggregate graph *A* obtained by compressing all the layers to a single graph. The measure used in De Domenico et al. [2015b] to differentiate the reduced multiplex from the aggregate graph *A* is termed as “relative entropy” and is given by

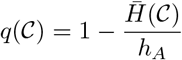

A larger value of *q*(C) is indicative of a more different from the aggregated network *A* and is suggestive of keeping the multiplex structure. As has been mentioned in De Domenico et al. [2015b], the goal is to find the optimal compression of layers by optimizing the relative entropy i.e. finding argmax [*q*(*C*)]. In this regard, they define the reducibility of the multiplex network *M* as follows

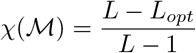

Here *χ*(*M*) = 0 would mean *L* = *L*_*opt*_ which means the multiplex is irreducible whereas *χ*(*M*) = 1 would mean *L*_*opt*_ = 1 suggesting that the aggregated graph is the most optimal choice. To overcome the complex problem of considering every single partition of the *L* layers, it is suggested in De Domenico et al. [2015b] to compute the pairwise Jensen-Shannon distance between layers to obtain a distance matrix, then cluster the layers hierarchically using this matrix, assess the resulting partitions with the relative entropy criterion *q*(·) and retain the partition with the highest *q*(·) i.e. the one most distinct from the aggregated network.

In a two-layer multiplex, we calculate the Von-Neumann entropy of the aggregated graph and the entropy per layer of the multiplex with two layers to evaluate the relative entropy which gives us the relative loss of information incurred if we used the aggregated network instead of the multiplex network.

### S4 Meso-scale Organization: Community Detection in Multiplex Networks

While a global measure of a network (typically a scalar-valued summary measure), gives an overall characterization of the network without specifically factoring in any measure of its internal organization, a meso-scale measure considers the organization of groups of nodes and is a measure to capture the intermediate structures in the multiplex such as communities, motifs, cores etc. in a network. We specifically look at the notion of communities in multiplex networks here.

Communities in multilayer networks are groups of nodes, formed within and across layers, that share similar characteristics when all the layers of interaction are taken into account. In a multiplex setting, each community can be viewed through its slices across the layers. In some cases, the same set of nodes form a particular community in every layer whereas in other instances the community’s membership varies from layer to layer due to addition or deletion of nodes in them. A useful perspective in this regard arises by noting that many real-world networks — whether social or biological — can be thought of as constraints on the flow of information, resources, or interactions, and that a multiplex network simply provides a richer description of those constraints. From this viewpoint, flow-based methods for community detection extend naturally to the multiplex setting as has been shown in De Domenico et al. [2015c].

To illustrate the idea, consider a social multiplex network in which nodes represent individuals and the layers represent different types of relationships, such as family, friendship, and workplace ties. Within each layer, restrictions on flow can lead to the emergence of modules, where interactions are more persistent and localized. Interestingly, these modules may or may not align across layers, as has been discussed earlier. For instance, if a group of friends jointly operate a business, the community detected in the friendship layer will reinforce the community detected in the work layer, effectively creating a unified module across both the layers. On the other hand, members of the same family may not necessarily overlap in friendships or professional contexts, so the family-based module may remain confined to its layer. Yet overlaps can still occur: a family member might also be a colleague or a close friend, thereby linking modules across layers.

In essence, communities in multiplex networks are modules formed due to sustained interaction among the nodes in them. This results in groups of nodes that trap information flow for long durations within their modules, while maintaining limited flow between them.

#### S4.1 Map Equation in Multiplex Networks

In the map equation framework, communities in a network are modules that capture the flow (communication/traffic) for longer stretches. This method uses the random walk as a proxy for this flow and identifies the modules in the network by compressing the description of this flow. De Domenico et al. [2015c] have generalized this concept to multilayer networks as “groups of nodes that capture flows within and across layers for a relatively long time”. It has been shown that the optimization of the description length of this flow leads to the identification of communities in the network i.e. this method seeks the node partition such that the expected description length of the flow is minimized. We give a brief overview of this proposed extension in the multiplex network context.

##### Representation

A multiplex network is a representation of a complex network which integrates the interactions of the nodes with each other differently through different means or at different points of time. These are often referred to as the different *modes* of interaction which represent the constraints on flow among the nodes in a given layer. Each physical node in a certain layer is termed as a ‘*state node*’. In the subsequent notations, layers are indexed by *l, m ∈*{1, 2, …, *L*} and the physical nodes are indexed by *i, j ∈*{1, 2, …, *N*}. A state node is denoted by (*i*, (*l*)).

##### Transition Probabilities when inter-layer links are known

In the presence of the empirical data on the inter-layer edges between the nodes, the transition probability from (*i*, (*l*)) to (*j*, (*m*)) is given by the following:

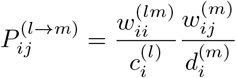

where 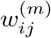 and 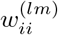 are the intra-layer edge weight from *i* to *j* in layer *m*, and the inter-layer replica coupling of node *i* between layers *l* and *m* respectively. 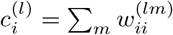 denote the inter-layer out strengths of node *i* in layer *l*, while 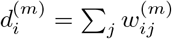 denotes the intra-layer out strengths of node *i* in layer *m*.

However, in many real life scenarios, there is inadequate data available on inter-layer links and that motivates the introduction of a relax rate *r* to model the inter-layer switches.

##### Transition Probabilities when inter-layer links are unknown

In such an instance, De Domenico et al. [2015c] propose to use a relax rate *r* to model the dynamics of the random walk across layers. In this case, the random walker moves only according to the intra-layer edges of a state node with probability 1 − *r* while this constraint is relaxed with probability *r* i.e. with probability *r*, the random walker can move along any edge of the physical node. Here, the transition probabilities are given by:

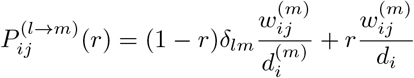

where 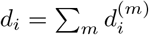

##### Stationary Distribution

The random walk is described with two sets of names, one for the modules, another set for the nodes inside the modules. The map equation optimizes the codelength i.e. the description of where the walker goes. The expected codelength of this description can be determined from the stationary distribution of the walk. The stationary distribution on the state nodes can be derived using the transition probabilities described earlier. Let 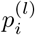 denote the stationary distribution of the state node (*i*, (*l*)) and it can be derived from

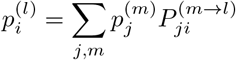

The stationary edge flow, per step, along the transition (*j*, (*m*)) to (*i*, (*l*)) is written as

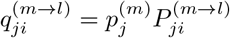

and hence we can write

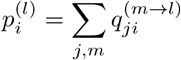

##### Multiplex Map Equation

Given a partition *P* that assigns each state node (*i*, (*l*)) to a module *k, k* = 1, 2, … *K*, we seek the partition that minimizes the expected description length *L*(*P*). The map equation framework encodes the random walk with one index codebook (for moves between modules) and one codebook per module (for visits within that module). By Shannon’s source-coding theorem, the average codeword length equals the entropy of the associated probability distribution of the relevant events, weighted by how often that codebook is used. Consequently, both the description lengths and their weights can be expressed in terms of the stationary visit rates 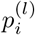 and the stationary edge flows 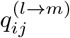.

For each module *k*, the entry and exit rates are defined as follows:

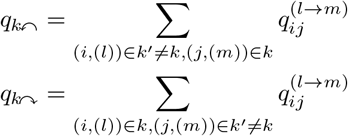

The module codebook for *k* contains one codeword for each physical node assigned to the module (aggregating all its state nodes in *k*) plus one exit codeword. The probability of the physical node *i* in *k* is

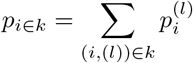

and the total use rate of the module codebook is 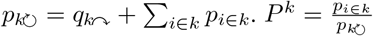 denotes the normalized probability distribution. The index codebook has one codeword per module, its total use rate is 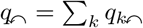 and the normalized probability distribution is 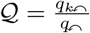

The map equation is the weighted sum of these entropies:

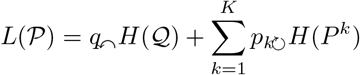

Minimizing *L*(*P*) favours modules with high persistence and low exit flow, i.e. groups that capture the random-walk flow for long stretches. In the multiplex setting, state nodes of the same physical node may appear in different modules and when multiple state nodes of a physical node fall in the same module, they share a single codeword.

##### Choice of the relax rate r

To choose the optimal value of the relax rate *r* in the multiplex Infomap analysis, we conducted certain simulation studies that reflect our belief of the underlying generative mechanism of the data i.e. the layers share a common mesoscale structure but are not identical. We simulated one hundred, two-layer multiplex networks from a stochastic block model type generative model in which the community labels were retained with a high probability and were randomly reassigned otherwise. This yielded moderately aligned communities across layers, with about 60 − 80% of nodes in the same community in both layers and the others had layer-specific allocations. To mimic the sparsity in the real data, the edges were drawn with low but relatively strong within-community connection probabilities and weaker between-community probabilities. We varied *r* in this setup and checked the recovery of the true community allocation measured using the variation of information metric. This metric was minimized around the values of *r* in the range [0.15, 0.3]. We therefore take the value of *r* as 0.15 for our analysis based on the optimal choice from our simulations and the recommendations in De Domenico et al. [2015c].

##### Summarization of communities in Multiplex Network

Our objective here is to assess the node level differences in the community allocations across the layers in the multiplex network and how this varies with age. To do so, for each of the *N* nodes in the multiplex network (let |*U* |= *N*), we define a *N ×* 1 layer-specific vector where nodes belonging to the same community take the value 1 and others take 0. Now, we compare these vectors for each node using the following metrics.

##### Variation of Information

This is an information theoretic approach proposed in Meilă [2007], for comparing clusters, by quantifying the change in the information due to switching from one cluster to the other. We discuss its intuitions briefly here. Let us suppose that there are *n* data points which have been clustered into *X* clusters *C*_*x*_, *x* = 1, 2, … *X*, where each cluster contains *n*_*x*_ data points. If we assume that picking each data point is equally likely, then it follows that the probability of a point landing in cluster *C*_*x*_ is

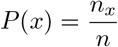

Thus with each clustering *C* is associated a discrete random variable taking *X* values and the uncertainty around each of these clustering can be quantified using the entropy of the clustering *C*

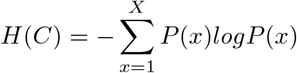

Let us denote by *P* (*x*), *x* = 1, 2 … *X* the random variables associated with clustering *C* and by *P* ^*′*^(*x*^*′*^), *x*^*′*^ = 1, 2 … *X*^*′*^ the random variables associated with clustering *C*^*′*^. Further, let *P* (*x, x*^*′*^) be the probability that a point belongs to cluster *C*_*x*_ in the clustering *C* and it belongs to *C*_*x*_*′* in the clustering *C*^*′*^. Then we have

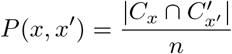

Then, the mutual information between the clusterings *C* and *C*^*′*^ is given by

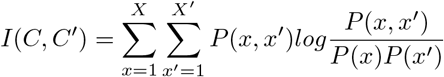

This form is also similar to the Kullback-Leibler divergence between the joint distribution and the product of the marginals and it gives a sense of the reduction in the uncertainty about *C*^*′*^ that comes with the knowledge of *C*. The mutual information cannot be greater than the total uncertainty in a clustering.

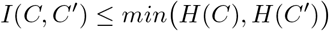

The variation of information between the clusterings *C* and *C*^*′*^ is given by

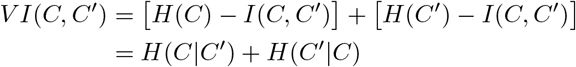

This is a sum of the conditional entropies quantifying how much uncertainty remains in each of the clusterings *C* and *C*^*′*^ after observing the other. This metric is bounded between 0 and *log*_2_(*n*) where 0 is indicative of identical partitions of the n elements while the upper bound is attained in case of independent partitions.

##### Jaccard Distance

Another metric used for the comparison of the community allocations for each node is the *Jaccard Distance Jaccard [1901], Levandowsky and Winter [1971]*. *We briefly discuss the idea here. Consider two finite sets A* and *B*, then the Jaccard Index is given by

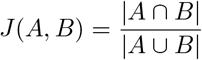

and the associated Jaccard distance is given by 1 − *J* (*A, B*). It follows from the above definition that 1 − *J* (*A, B*) ∈ (0, 1). The closer the value of the distance to 1, the more dissimilar the two sets are.

## References

Wan He, Daniel Goodkind, Paul R Kowal, et al. An aging world: 2015, volume 2023. United States Census Bureau Washington, DC, 2016.

Chun-Yi Zac Lo, Yong He, and Ching-Po Lin. Graph theoretical analysis of human brain structural networks. 2011.

Ke Liu, Shixiu Yao, Kewei Chen, Jiacai Zhang, Li Yao, Ke Li, Zhen Jin, and Xiaojuan Guo. Structural brain network changes across the adult lifespan. Frontiers in aging neuroscience, 9:275, 2017.

Josh Neudorf, Kelly Shen, and Anthony R McIntosh. Reorganization of structural connectivity in the brain supports preservation of cognitive ability in healthy aging. Network Neuroscience, 8(3):837–859, 2024.

Manon Edde, Gaëlle Leroux, Ellemarije Altena, and Sandra Chanraud. Functional brain connectivity changes across the human life span: From fetal development to old age. Journal of neuroscience research, 99(1):236–262, 2021.

Sandra Doval, Alberto Nebreda, and Ricardo Bruña. Functional connectivity across the lifespan: a cross-sectional analysis of changes. Cerebral Cortex, 34(10):bhae396, 2024.

Alexa Mousley, Richard AI Bethlehem, Fang-Cheng Yeh, and Duncan E Astle. Topological turning points across the human lifespan. Nature communications, 16(1):10055, 2025.

Micaela Y Chan, Denise C Park, Neil K Savalia, Steven E Petersen, and Gagan S Wig. Decreased segregation of brain systems across the healthy adult lifespan. Proceedings of the National Academy of Sciences, 111(46):E4997–E5006, 2014.

Lubin Wang, Longfei Su, Hui Shen, and Dewen Hu. Decoding lifespan changes of the human brain using resting-state functional connectivity mri. 2012.

Shania Mereen Soman, Nandita Vijayakumar, Phoebe Thomson, Gareth Ball, Christian Hyde, and Timothy J Silk. Cortical structural and functional coupling during development and implications for attention deficit hyperactivity disorder. Translational psychiatry, 13(1):252, 2023.

Graham L Baum, Zaixu Cui, David R Roalf, Rastko Ciric, Richard F Betzel, Bart Larsen, Matthew Cieslak, Philip A Cook, Cedric H Xia, Tyler M Moore, et al. Development of structure–function coupling in human brain networks during youth. Proceedings of the National Academy of Sciences, 117(1):771–778, 2020.

Guozheng Feng, Yiwen Wang, Weijie Huang, Haojie Chen, Jian Cheng, and Ni Shu. Spatial and temporal pattern of structure–function coupling of human brain connectome with development. Elife, 13:RP93325, 2024.

Meredith A Shafto, Lorraine K Tyler, Marie Dixon, Jason R Taylor, James B Rowe, Rhodri Cusack, Andrew J Calder, William D Marslen-Wilson, John Duncan, Tim Dalgleish, et al. The cambridge centre for ageing and neuroscience (cam-can) study protocol: a cross-sectional, lifespan, multidisciplinary examination of healthy cognitive ageing. BMC neurology, 14(1):204, 2014.

Jason R Taylor, Nitin Williams, Rhodri Cusack, Tibor Auer, Meredith A Shafto, Marie Dixon, Lorraine K Tyler, Richard N Henson, et al. The cambridge centre for ageing and neuroscience (cam-can) data repository: Structural and functional mri, meg, and cognitive data from a cross-sectional adult lifespan sample. neuroimage, 144:262–269, 2017.

Manlio De Domenico, Mason A Porter, and Alex Arenas. Muxviz: a tool for multilayer analysis and visualization of networks. Journal of Complex Networks, 3(2):159–176, 2015a.

Manlio De Domenico. Multilayer networks: Analysis and visualization. Introduction to muxViz with R. Cham: Springer, 2022.

Federico Battiston, Vincenzo Nicosia, Mario Chavez, and Vito Latora. Multilayer motif analysis of brain networks. Chaos: An Interdisciplinary Journal of Nonlinear Science, 27(4), 2017.

Federico Battiston, Jeremy Guillon, Mario Chavez, Vito Latora, and Fabrizio De Vico Fallani. Multiplex core–periphery organization of the human connectome. Journal of the Royal Society Interface, 15(146), 2018.

Sol Lim, Filippo Radicchi, Martijn P van den Heuvel, and Olaf Sporns. Discordant attributes of structural and functional brain connectivity in a two-layer multiplex network. Scientific reports, 9(1):2885, 2019.

Maria Grazia Puxeddu, Joshua Faskowitz, Olaf Sporns, Laura Astolfi, and Richard F Betzel. Multi-modal and multi-subject modular organization of human brain networks. NeuroImage, 264:119673, 2022.

Nicholas Parsons, Julien Ugon, Kerri Morgan, Sergiy Shelyag, Alex Hocking, Su Yuan Chan, Govinda Poudel, Karen Caeyenberghs, et al. Structural-functional connectivity bandwidth of the human brain. NeuroImage, 263:119659, 2022.

Gwendolyn Jauny, Mite Mijalkov, Anna Canal-Garcia, Giovanni Volpe, Joana Pereira, Francis Eustache, and Thomas Hinault. Linking structural and functional changes during aging using multilayer brain network analysis. Communications biology, 7(1):239, 2024.

Martijn Devrome, Koen Van Laere, and Michel Koole. Multiplex core of the human brain using structural, functional and metabolic connectivity derived from hybrid pet-mr imaging. Frontiers in neuroimaging, 2:1115965, 2023.

Maedeh Khalilian, Monica N Toba, Martine Roussel, Sophie Tasseel-Ponche, Olivier Godefroy, and Ardalan Aarabi. Age-related differences in structural and resting-state functional brain network organization across the adult lifespan: A cross-sectional study. Aging Brain, 5:100105, 2024.

Christopher H van Dyck, John P Seibyl, Robert T Malison, Marc Laruelle, Sami S Zoghbi, Ronald M Baldwin, and Robert B Innis. Age-related decline in dopamine transporters: analysis of striatal subregions, nonlinear effects, and hemispheric asymmetries. The American journal of geriatric psychiatry, 10(1):36–43, 2002.

Jean-Claude Dreher, Andreas Meyer-Lindenberg, Philip Kohn, and Karen Faith Berman. Age-related changes in midbrain dopaminergic regulation of the human reward system. Proceedings of the National Academy of Sciences, 105(39):15106–15111, 2008.

Timothy A Salthouse. The processing-speed theory of adult age differences in cognition. Psychological review, 103(3): 403, 1996.

Denise C Park and Patricia Reuter-Lorenz. The adaptive brain: aging and neurocognitive scaffolding. Annual review of psychology, 60:173–196, 2009.

Kang Min Park, Keun Tae Kim, Dong Ah Lee, Gholam K Motamedi, and Yong Won Cho. Structural and functional multilayer network analysis in restless legs syndrome patients. Journal of Sleep Research, 33(3):e14104, 2024.

Dandan Li, Yating Zhang, Luyao Lai, Jianchao Hao, Xuedong Wang, Zhenyu Zhao, Xiaohong Cui, Jie Xiang, and Bin Wang. The impact of indirect structure on functional connectivity in schizophrenia using a multiplex brain network. Journal of Psychiatric Research, 179:257–265, 2024.

Juan F Quinones, Carsten Gießing, Axel Heep, and Andrea Hildebrandt. Audiovisual integration and whole-brain networks in preterm and full-term neonates: A two-layer multiplex network perspective on structural and functional connectivity. Imaging Neuroscience, 3:IMAG–a, 2025.

Matthew F Glasser, Timothy S Coalson, Emma C Robinson, Carl D Hacker, John Harwell, Essa Yacoub, Kamil Ugurbil, Jesper Andersson, Christian F Beckmann, Mark Jenkinson, et al. A multi-modal parcellation of human cerebral cortex. Nature, 536(7615):171–178, 2016.

Bruce Fischl. Freesurfer. Neuroimage, 62(2):774–781, 2012.

J-Donald Tournier, Robert Smith, David Raffelt, Rami Tabbara, Thijs Dhollander, Maximilian Pietsch, Daan Christiaens, Ben Jeurissen, Chun-Hung Yeh, and Alan Connelly. Mrtrix3: A fast, flexible and open software framework for medical image processing and visualisation. Neuroimage, 202:116137, 2019.

Mikko Kivelä, Alex Arenas, Marc Barthelemy, James P Gleeson, Yamir Moreno, and Mason A Porter. Multilayer networks. Journal of complex networks, 2(3):203–271, 2014.

Manlio De Domenico, Albert Solé-Ribalta, Emanuele Cozzo, Mikko Kivelä, Yamir Moreno, Mason A Porter, Sergio Gómez, and Alex Arenas. Mathematical formulation of multilayer networks. Physical Review X, 3(4):041022, 2013.

Stefano Boccaletti, Ginestra Bianconi, Regino Criado, Charo I Del Genio, Jesús Gómez-Gardenes, Miguel Romance, Irene Sendina-Nadal, Zhen Wang, and Massimiliano Zanin. The structure and dynamics of multilayer networks. Physics reports, 544(1):1–122, 2014.

Manlio De Domenico, Vincenzo Nicosia, Alexandre Arenas, and Vito Latora. Structural reducibility of multilayer networks. Nature communications, 6(1):6864, 2015b.

Manlio De Domenico, Andrea Lancichinetti, Alex Arenas, and Martin Rosvall. Identifying modular flows on multilayer networks reveals highly overlapping organization in interconnected systems. Physical Review X, 5(1):011027, 2015c.

Marina Meilă. Comparing clusterings—an information based distance. Journal of multivariate analysis, 98(5):873–895, 2007.

Paul Jaccard. Étude comparative de la distribution florale dans une portion des alpes et des jura. Bull Soc Vaudoise Sci Nat, 37:547–579, 1901.

Michael Levandowsky and David Winter. Distance between sets. Nature, 234(5323):34–35, 1971.

Mark Newman. Networks. Oxford university press, 2018.

